# Reconsidering electrophysiological markers of response inhibition in light of trigger failures in the stop-signal task

**DOI:** 10.1101/658336

**Authors:** P Skippen, W. R Fulham, P.T Michie, D Matzke, A Heathcote, F Karayanidis

## Abstract

We investigate the neural correlates underpinning response inhibition using a parametric ex-Gaussian model of stop-signal task performance, fit with hierarchical Bayesian methods, in a large healthy sample (N=156). The parametric model accounted for trigger failure (i.e., failures to initiate the inhibition process) and returned an SSRT estimate (SSRT_EXG3_) that was attenuated by ≈65ms compared to traditional non-parametric SSRT estimates (SSRT_int_). The amplitude and latency of the N1 and P3 event related potential components were derived for both stop-success and stop-failure trials and compared to behavioural estimates derived from traditional (SSRT_int_) and parametric (SSRT_EXG3_, trigger failure) models. Both the fronto-central N1 and P3 peaked earlier and were larger for stop-success than stop-failure trials. For stop-failure trials only, N1 peak latency correlated with both SSRT estimates as well as trigger failure and temporally coincided with SSRT_EXG3_, but not SSRT_int_. In contrast, P3 peak and onset latency were not associated with any behavioural estimates of inhibition for either trial type. While overall the N1 peaked earlier for stop-success than stop-failure trials, this effect was not found in poor task performers (i.e., high trigger failure/slow SSRT). These findings are consistent with attentional modulation of both the speed and reliability of the inhibition process, but not for poor performers. Together with the absence of any P3 onset latency effect, our findings suggest that attentional mechanisms are important in supporting speeded and reliable inhibition processes required in the stop-signal task.

## 1. Introduction

Response inhibition is a core component of cognitive control that is associated with the cancelation or suppression of an inappropriate behaviour (Bari & Robbins, 2013) and is typically operationalised using the stop-signal task (Logan & Cowan, 1984). In the stop-signal task, participants engage in a speeded choice response task. On a small proportion of randomly selected trials, a stop signal occurs after the choice stimulus and participants must inhibit their prepared response. The prominent horse-race model of the stop-signal task estimates the latency of the ‘stop process’ or stop-signal reaction time (SSRT), which is assumed to fully characterise stop-signal task performance, without the need to account for other processes, such as attention.

Increased SSRT in people with attention deficit hyperactivity disorder, schizophrenia, and substance use disorders is thought to indicate less efficient response inhibition (for a review, see Lipszyc & Schachar, 2010), and to result in reduced impulse control in healthy cohorts (Sharma, Markon, & Clark, 2014; Skippen et al., 2019). However, the purity of SSRT as *the* sole measure of inhibitory control has been questioned (Band, Van Der Molen, & Logan, 2003; Chatham, et al., 2012; Logan, 1994; Matzke, Curley, Gong, & Heathcote, 2019; Matzke, Hughes, Badcock, Michie, & Heathcote, 2017; Matzke, Love, & Heathcote, 2017; Skippen et al., 2019; Verbruggen, Best, Bowditch, Stevens, & McLaren, 2014; Verbruggen, McLaren, & Chambers, 2014).

### 1.1. Implications of biased SSRT estimations

The horse-race model (Logan & Cowan, 1984) attributes the outcome of any given stop trial to a race between two independent processes: the go process (measured by go trial RT) and the stop process (measured by SSRT). If, on a given trial, the go process finalises before the stop process, the response will not be inhibited (i.e., stop-failure trial). Alternatively, if the stop process finalises before the go process, the response will be successfully inhibited (i.e., stop-success trial). However, if we assume that, on a given trial, either the go or the stop process fails to initiate, the estimated SSRT is no longer a pure measure of response inhibition. If the stop process is not triggered, the response to the go trial cannot be inhibited. Likewise, if the go process is not initiated, the stop process can be extremely slow yet still result in successful inhibition. Traditional, non-parametric estimation techniques^1^, such as the mean or integration method (see Verbruggen et al., 2019) do not account for failures to engage the go or the stop process, and therefore may result in biased estimates of SSRT (Band et al., 2003; Matzke, Curley, et al., 2019; Matzke, Love, et al., 2017; Skippen et al., 2019). While adjustments to non-parametric methods have been developed to account for omission rates on go trials, as a close proxy for go failures (e.g., Tannock, Schachar, Carr, Chajczyk, & Logan, 1989), simulations suggest that this method still over-estimates SSRT (Verbruggen et al., 2019). The method with the least bias involves replacing omissions on go trials with the slowest go RT for each participant (e.g., Verbruggen et al., 2019).

Trigger failure, or the failure to initiate the stop process, has long been acknowledged as possibly contributing to performance on the stop-signal task (Logan, 1994), but cannot easily be taken into account with non-parametric methods of estimating SSRT (Band et al., 2003). Not accounting for trigger failure has been shown to produce biased estimates of SSRT (Band et al., 2003; Jana, Hannah, Muralidharan, & Aron, 2020; Matzke, Love, et al., 2017; Skippen et al., 2019; Weigard, Heathcote, Matzke, & Huang-Pollock, 2019), indicating that effective inhibitory control relies not only on fast SSRT, but also on the reliability of triggering the stop process (Chatham et al., 2012; Matzke, Love, et al., 2017).

We have previously described a parametric model of the stop-signal task in which the finishing times of the stop and go processes follow an ex-Gaussian distribution, and model parameters are estimated with the Bayesian Estimation of ex-Gaussian Stop-Signal Reaction Time Distribution (BEESTS) procedure (Matzke, Dolan, Logan, Brown, & Wagenmakers, 2013; Matzke, Love, et al., 2013). This model estimates the entire distribution of SSRT rather than a single summary measure (see Matzke, Verbruggen, & Logan, 2018, for an overview of the different estimation methods). In this study, we use an ex-Gaussian model (EXG3; Matzke, Curley, et al., 2019) that extends the BEESTS model by incorporating failure to trigger a response to the go (go failure) and the stop (trigger failure; see also Matzke, Love, et al., 2017) stimulus, as well as accounting for go errors through separate racers that correspond to correct (matching) and error (mis-matching) responses to the go stimulus.

The practical importance of modelling trigger and go failures has been highlighted in a number of studies (Band et al., 2003; Matzke, Curley, et al., 2019; Matzke, Dolan, et al., 2013; Matzke, Hughes, et al., 2017; Skippen et al., 2019; Weigard, Heathcote, Matzke, & Huang-Pollock, 2019). For example, Skippen et al. (2019) found that, in healthy young participants, the SSRT estimate derived from the EXG3 model was ≈100ms faster than the estimate obtained using the traditional non-parametric integration method (e.g., Tannock et al., 1989). Critically, this changed the relationships between SSRT and measures of impulsivity. Using the non-parametric integration technique, impulsivity was correlated with estimated SSRT. However, using estimates from the EXG3 model, impulsivity was correlated with trigger failure and go failure rate but not SSRT. Using a similar ex-Gaussian model, Matzke, Hughes et al. (2017) showed that people with schizophrenia had both higher levels of trigger failure and slower SSRT when compared to controls^2^. The distribution of SSRTs suggested that the delay in the stop process was largely due to poor encoding of the stop-signal, rather than a slower inhibition process *per se*. The increased probability of trigger failure and the delayed initiation of the stop process in people with schizophrenia was interpreted as indicative of an attention deficit. Therefore, attentional factors may play a larger role in inhibitory ability than previously thought. In this study, we take a model-based neuroscience approach (Boucher, Palmeri, Logan, & Schall, 2007; Wiecki & Frank, 2013) to characterise the mechanisms that contribute to the stop process and trigger failure (Sebastian, Forstmann, & Matzke, 2018), using electroencephalogram (EEG) activity recorded during the stop-signal task to capture the timeline of cognitive processes involved in response inhibition.

### 1.2. Temporal processes of response inhibition

The auditory stop signal typically elicits a number of event-related potential (ERP) components commonly measured over the fronto-central scalp, including an early N1 (≈100-200 ms post-event and a later P3 (≈300-350 ms; Pires, Leitão, Guerrini, & Simões, 2014). Visual stop signals also generate a frontal N2 component (≈200 ms) that is not typically evident with auditory stop signals (see Kenemans, 2015).

The N1 is thought to reflect the level of stimulus encoding in the auditory cortex (Hillyard, Hink, Schwent, & Picton, 1971; Luck, Woodman, & Vogel, 2000; Näätänen & Picton, 1987). It is comprised of several underlying components that are influenced by voluntary attention to the stimulus, the physical attributes of the stimulus, and the conditions under which it is presented (Näätänen, 1982; Näätänen, Gaillard, & Mäntysalo, 1978; Näätänen & Michie, 1979; Näätänen & Picton, 1987). In the response inhibition literature, N1 is commonly used as a measure of the impact of the stop signal on auditory cortex processing, which may vary depending on fluctuations in attention (Bekker, Kenemans, Hoeksma, Talsma, & Verbaten, 2005; Dimoska & Johnstone, 2008). The commonly reported ‘stop-N1’ effect (i.e., increased N1 amplitude for stop-success compared to stop-failure trials) suggests that N1 amplitude is associated with the outcome of the stop process (Bekker, Kenemans, et al., 2005; Hughes, Fulham, Johnston, & Michie, 2012; Lansbergen, Böcker, Bekker, & Kenemans, 2007). This ‘stop-N1’ effect is suggested to reflect a mechanism that primes the detection of the stop-signal, in turn potentiating a faster connection between brain regions vital to improving the speed of inhibition (Kenemans, 2015).

The role of the later P3 component has been the subject of substantial debate. Some argue that the P3 represents the manifestation of the inhibition process (for a review, see Huster, Enriquez-Geppert, Lavallee, Falkenstein, & Herrmann, 2013) and this is based largely on the finding that P3 latency tends to coincide with SSRT latency (Kok, Ramautar, De Ruiter, Band, & Ridderinkhof, 2004; Wessel & Aron, 2015). However, others have argued that P3 peaks too late to represent inhibition and is more likely to reflect post-inhibition processing (González-Villar, Bonilla, & Carrillo-de-la-Peña, 2016; Huster et al., 2013; Ramautar, Kok, & Ridderinkhof, 2004). Importantly, both arguments are based on conventional estimates of SSRT, which as discussed above, may be biased. Given that models that account for trigger failure produce attenuated estimates of SSRT, the temporal association between stop-ERPs (especially the P3) and SSRT is likely to be altered. For example, SSRT estimates derived from the EXG3 model in Skippen et al (2019) aligned the end of the stop process within the latency range of N1, and at least 100ms before the onset of the typical P3. Therefore, models that include trigger failure in the estimation of SSRT may challenge previous interpretations of the ERP correlates of response inhibition.

Matzke, Hughes et al. (2017) is the only study to investigate the relationship between ERP components and parameters from an ex-Gaussian model of the stop-signal task. An earlier stop-N1 latency was associated with a higher probability of trigger failure, in a small group of people with schizophrenia. At face value, this relationship is counterintuitive, suggesting that earlier processing of the stop signal (as indexed by the N1) is predictive of less efficient inhibition (i.e., higher rate of trigger failure). However, as the N1 is comprised of a number of partially overlapping sub-components (Näätänen & Picton, 1987), an earlier N1 peak latency may result from attenuation of a late N1 sub-component and may not necessarily represent faster processing of the stop signal (Matzke, Hughes et al., 2017). This would be consistent with the hypothesis that trigger failure results from the dysfunctional encoding of the attributes of the stop signal, which then fails to generate the appropriate response pattern (i.e., inhibition). However, these findings require replication as they were based on a small group (*N* ≤ 13 of patients) and did not differentiate between stop-success and stop-failure trials.

### 1.3. Current study

In the present study, we examine the relationship between EXG3 parameters and stop-signal ERP components in a large, young, healthy cohort (*N* = 156). With high statistical power and opportunity to examine individual variability, we aim to provide insight into the neural processes that underlie trigger failure, go failure, and SSRT. Based on Matzke, Hughes et al. (2017), we expect that trigger failure rate will be negatively associated with stop-N1 peak latency. We also estimate N1 onset latency to examine whether rate of trigger failure is associated with a relative enhancement of the early N1 sub-component. In line with the attentional account of trigger failure, we expect that lower levels of attention to the stop-signal (i.e., reduced N1 amplitude) will be associated with higher rates of trigger failure. Moreover, we examine whether EXG3 model parameters are differentially associated with the success or failure of the inhibition process, by deriving ERP components for both successful and failed stop trials.

As SSRT latency is attenuated after accounting for trigger failure, we expect that the relationship between SSRT and ERP components commonly reported in the literature will be altered. Specifically, we expect to challenge the common interpretation of the P3 latency as an index of the response inhibition process by showing that, when accounting for trigger failure, the relationship between SSRT and P3 latency will be weak. We compare the relationship of onset and peak P3 latency with both traditional and EXG3 estimates of SSRT. Given recent suggestions that response inhibition may be a more automatic process than previously thought (Verbruggen, McLaren, & Chambers, 2014), we expect that SSRT estimated using the EXG3 model will be more strongly associated with the earlier N1 than SSRT derived using traditional estimation. Finally, we run exploratory analyses of the relationship between other EXG3 model parameters (i.e., go RT, go failures) and ERP component measures to inform future hypotheses about the neural correlates of response inhibition in a healthy young sample.

## 2. Methods

### 2.1. Participants and procedure

The data reported here were collected as part of a larger longitudinal study, the Age-ility Project (Karayanidis et al., 2016). The sample overlaps with that reported by Skippen et al. (2019), with the additional exclusion of participants without clean EEG. After loss to attrition between testing sessions (see Karayanidis et al., 2016), 208 participants attempted the stop-signal task, but the final sample included data from 156 participants (see *2.3 Data Analysis* for all exclusion criteria). This study conforms to the Declaration of Helsinki and was approved by the University of Newcastle Human Research Ethics Committee (HREC: H-2012-0157).

### 2.2. Stimuli and apparatus

#### 2.2.1. Stop-signal task

The primary go task was a two-choice number parity task (700 trials). The stimulus was a number between 2 and 9, presented for 100ms in the centre of a grey rectangle. On 29% of trials (≈200 trials) the go stimulus was followed, after a variable stop-signal delay, by an auditory stop signal that was delivered binaurally through calibrated headphones (1000Hz, 85dB tone, 100ms duration). The stop-signal delay ranged from 50-800ms and decreased or increased by 50ms after every failed or successful stop trial, respectively. Following a single practice block, behavioural responses and EEG activity were recorded for 700 trials across five blocks.

#### 2.2.2. Electroencephalogram recording

EEG data were recorded (2048Hz sample rate, bandpass filter of DC-400Hz) via a BioSemi Active Two system with 64 scalp electrodes as well as bilateral mastoid, bilateral ocular, and infra/supra ocular sites. Common mode sense and driven right leg electrodes were positioned inferior to P1 and P2, respectively. Data were recorded relative to an amplifier reference voltage and were re-referenced to the common average offline to remove common-mode signals. See *2.3.3*, for EEG processing details.

### 2.3. Data analysis

#### 2.3.1. Data Cleaning

A technical issue resulted in the exclusion of the first 20 participants. Another 32 participants were excluded due to poor or non-compliant performance on the stop-signal task. Of these: (a) Nine participants slowed their go trial reaction time by more than 300ms over the course of the experiment*^3^*, most likely to facilitate inhibition on a subsequent stop trial. This slowing of responses can bias SSRT estimates; (b) Four participants responded on over 75% of stop trials, in violation of what would be expected given the stop-signal delay algorithm; (c) Sixteen participants violated the independence assumption of the horse race model. This assumption states that the go and stop processes are independent and is tested by confirming that mean RT on stop trials (i.e., stop-failure trials) is not slower than mean RT on go trials (Logan & Cowan, 1984; Verbruggen et al., 2019; (d) One violated both independence and response rate assumptions; (e) One had both independence violations and commission error rates on the go task approaching chance; and (f) One had no errors on the go task and therefore could not be modelled with EXG3. This resulted in an exclusion rate of approximately 16.5% (not including the technical issue).

Six participants had a block of behavioural data that differed from the rest of their performance. Of these: One participant made no responses with their right hand across the entire 5th block; One reported forgetting to stop to the signal in the first block but corrected this for the remaining blocks; Three made no responses at all in one block (1^st^, 2^nd^, 4^th^, respectively); and, One participant made no successful stops in the first block. For each participant, the aberrant block was removed. The stop-signal delay tracking recovered quickly in the following block of trials for all participants.

#### 2.3.2. Modelling of Response Inhibition

The EXG3 model (Matzke, Curley, et al.. 2019) assumes that finishing times of the three runners (i.e., two go runners and one stop runner) can be described by an ex-Gaussian distribution. It estimates the finishing time of both the matching (correct) and mismatching (error) response to the go stimuli. For each go process and the stop process, the ex-Gaussian distribution has three parameters μ, σ, and τ, which characterise the mean and standard deviation of the normal component, and the mean of the exponential component (i.e., the long slow tail of the distribution), respectively. The mean and variance of each finishing time distribution can then be estimated as μ + τ and σ^2^ + τ^2^, respectively. In order for the probability of go and trigger failure parameters to be modelled with a normal group-level distribution, they are first projected from the probability scale to the real line with a “probit” (i.e., standard normal cumulative distribution function) transformation (see also, Matzke, Dolan, Batchelder, & Wagenmakers, 2015; Rouder, Lu, Morey, Sun, & Speckman, 2008). After modelling is complete, they are returned to the probability scale using a bivariate inverse probit transformation for further analysis and interpretation.

We used the Dynamic Models of Choice (DMC; Heathcote et al., 2019) software implemented in R (R Core Team, 2018) to estimate model parameters via Bayesian hierarchical modelling. In hierarchical modelling, the population-level mean and standard deviation parameters characterise the population-level distribution for each model parameter. Weakly informative uniform priors were set for the population-level parameters, which are identical to those of Skippen et al. (2019). Posterior distributions of the parameters were obtained using Differential Evolution Markov Chain Monte Carlo (MCMC) sampling (Ter Braak, 2006), with steps closely mimicking Heathcote et al. (2019).

To confirm the non-negligible presence of trigger failures, we ran two EXG3 models, one with a trigger failure parameter and one without. We ran 33 MCMC chains in the model with trigger failure and 30 in the model without (e.g., three times as many chains as model parameters). Participants were initially modelled separately until the MCMC chains reached convergence, with thinning of every 10^th^ sample. These individual fits were then used as the start values for the hierarchical fits. First, we set a 5% probability of migration steps replacing cross-over steps for both the participant and the hierarchical levels and sampled until there were no outlying chains. Then, pure crossover steps were performed until chains were converged and stable. After this, an additional 200 samples per chain were retained as the final set from which further analysis is undertaken. Convergence was confirmed by visual inspection and Gelman-Rubin 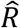 (Gelman & Rubin, 1992) values below 1.1. Effective sample size was checked to ensure that all parameters were well characterized by posterior distributions.

The population distributions describe the between-subject variability of the parameters and are appropriate for population inference, analogous to frequentist random-effects analysis. The individual participant parameters are useful for examining individual differences and the relationship between model parameters and ERP measures.

#### 2.3.3. Electrophysiological Data

EEG data pre-processing was performed using in-house techniques within EEGDisplay v6.4.4 (Fulham, 2015). Firstly, the continuous EEG data were downsampled to 512Hz and visually inspected to exclude intervals containing gross artefact. Bad EEG channels were replaced by interpolating adjacent scalp electrodes. Eye blink artefact was corrected using a regression-based procedure (Semlitsch, Anderer, Schuster, & Presslich, 1986). As a secondary control measure to visual inspection, gross movement or muscle artefacts were excluded with an automated procedure which applies amplitude thresholds within specific frequency bands. To control for residual eye movement artefact, we applied a threshold four times the standard deviation (SD) of signal < 2Hz. To target muscular artefact, a threshold of 7.5 times the SD in signal > 10Hz was used. To control for any other residual artefact, a threshold of 6 times the SD was set for signal between 0.5-15Hz. Finally, the signal was bandpass filtered between 0.05 and 30Hz. Participants had an average of 1.05 channels interpolated (± SD = 2.24), none of which were FCz. An average of 162 (± 24) stop trials (from 200) remained after artefact corrections.

EEG epochs for the stop-signal trials were extracted from 900ms pre-event to 1400ms post-event and averaged separately for stop-success and stop-failure trials. As stop-locked ERPs are contaminated by the overlapping ERP response to the preceding go stimulus (especially for short stop-signal delays), we extracted the stop-locked ERPs by applying level-1 ADJAR correction, based on the assumption that the responses elicited by go and stop stimuli combine linearly. This procedure takes the convolution of the go-locked ERP with the stop-signal delay distribution, and subtracts this from the stop-locked ERP (see Woldorff, 1993)^4^. To remove any remaining peri-event baseline shifts that may affect amplitude measures and to control for any residual activity from the previous Go stimulus, we baseline corrected using a 50ms peri-stop baseline (i.e., −25 pre-to 25ms post-stop).

Before ERP component measurement, we re-referenced the data using a surface Laplacian transformation (or current source density, Kayser & Tenke, 2006, 2015; Nunez & Srinivasan, 2006) which we have previously shown in this sample to differentiate between partially overlapping components (Wong et al., 2018). The surface Laplacian is insensitive to broad changes in signal, resulting from volume conduction and reference choice and is more sensitive to activity from cortical generators, resulting in improved spatial and temporal information (for a technical review, see Hjorth, 1975; Yao, 2002; Kayser & Tenke, 2015). A spherical spline function was applied using the CSD toolbox in MATLAB across all scalp electrode locations, with the spline flexibility parameter *m* = 4, for increased rigidness (Kayser & Tenke, 2006, 2015). As the EEG signal is transformed based on the second partial derivative of the signal (μV) over a spatial area (cm^2^–i.e., the scalp), the measurement scale is μV/cm^2^.

Component selection targeted two fronto-central components typically elicited by auditory stop signals (N1, P3; Huster et al., 2013; Kok, Ramautar, De Ruiter, Band, & Ridderinkhof, 2004). To increase the resolution of latency estimates, we interpolated waveforms by a factor of four to a 2048Hz sampling rate before estimating component measures. Peak latency was estimated using fractional area latency across a specified window after stop-signal onset (N1 = 80 – 180ms; P3 = 180 – 350ms; Kiesel, Miller, Jolicœur, & Brisson, 2008). N1 and P3 peak amplitude was measured using an average of 20ms around the respective peak latency value for each individual. The onset latency of each component was estimated by determining the latency at which the amplitude reached 50% of the component’s peak amplitude. Based on previous literature, we expected that the midline frontal/fronto-central sites would show the largest effects for stop-related ERP components. Head topographies, as well as trial type differences (described in 2.4.3) were used to determine FCz as the site of interest.

#### 2.3.4. Plausible Values Analysis

Traditional tests of correlations between hierarchical model parameters and ERP component measures ignore both the uncertainty of the parameter estimates and the effects of hierarchical shrinkage, and therefore tend to be overconfident. We avoid these problems by using a plausible value analysis (Ly et al., 2018) to evaluate these relationships. This calculates a distribution of correlations between covariates (e.g., N1 latency) and model parameters (i.e., mean SSRT) using each MCMC sample from the posterior distribution of the parameter. This process results in a set of ‘plausible’ values of the sample correlation, *r*, for *n* individuals. As described by Ly et al., the sample correlation can be transformed into the posterior distribution of the population correlation (ρ), which is a function of *r* and *n*. Repeating this process for all plausible sample correlations and pointwise averaging the population correlation distribution yields the estimated posterior distribution of the population correlation.

### 2.4. Statistical Procedures

#### 2.4.1. Stop-Signal Task Behaviour

We report task behaviour using the Verbruggen et al. (2019) recommendations. The traditional SSRT estimate, SSRT_int_, was obtained using the integration technique described in Verbruggen et al. (2019), where go omissions trials are replaced with the participant’s maximum go RT.

#### 2.4.2. Model Selection and Parameter Estimation

To examine whether the presence of trigger failures was non-negligible, we compared hierarchical models with and without trigger failure using the Deviance Information Criterion (DIC; Spiegelhalter, Best, Carlin, & Van Der Linde, 2002). A difference in DIC of 10 or more is taken as substantial evidence in favour of the model with the smaller DIC. As set out in Heathcote et al (2019), we assessed the absolute goodness-of-fit using posterior predictive model checks (Gelman, Meng, & Stern, 1996). We used the most supported EXG3 model to estimate mean SSRT (i.e., μ_stop_ + τ_stop_), mean finishing time of the matching go runner (i.e., μ_go correct_ + τ_go correct_), go failures, and trigger failures. We refer to the EXG3-based estimate of mean SSRT as SSR_TEXG3_.

#### 2.4.3. ERP Component Electrode Site/s of Interest

We compared the scalp topography of the grand average ERP components with previous literature, and used a Bayesian paired t-test to determine the site with the largest amplitude differences between trial types (i.e., stop-failure vs stop-success) for each ERP component. Two-sided Bayesian t-tests were computed using the *BayesFactor* package (Morey & Rouder, 2018) in R (R Core Team, 2018), with effect sizes given as the posterior median effect size (δ). Default “medium” Cauchy priors with 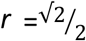 were used as priors for the t-test on trial type. Based on these results, and the head topographies, we selected FCz for measurement of N1 and P3 components.

#### 2.4.4. Traditional Correlational Analysis

Correlations between SSRT_int_ and ERP component measures were undertaken using Pearson product moment correlations. Inference was conducted using Bayes factors as estimated in the *BayesFactor* R package (Morey & Rouder, 2018; R Core Team, 2018). Correlation tests were completed with a prior on ρ centred around zero, with a default “medium” *r* scale argument of 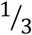. Kass and Raftery (1995) conventions describe the strength of evidence that the data provide for the alternative hypothesis. For the null and alternative, a Bayes factor between 1/3 and 3 is considered ‘equivocal’ (i.e., indicating more data are needed to obtain a clear outcome), between 1/3-1/20 or 3-20 is considered ‘positive’, between 1/20-1/150 or 20-150 ‘strong’, and Bayes factors less than 1/150 or greater than 150 are considered ‘very strong’.

#### 2.4.5. Plausible Values Analysis

To identify overlap between traditional and model estimates of response inhibition, we examined the relationship between the posterior distributions of the mean SSRT_EXG3_, trigger failure and go failure parameters, and SSRT_int_ using plausible values analysis. Default priors for the plausible values analysis provide a uniform distribution, with equal likelihood for all values between −1 and 1. We also report plausible value correlations between model parameters and ERP component measures from both stop-success and stop-failure trials. All plausible values relationships are accompanied by associated Bayesian *p*-values, which denote the proportion of the distribution that is shifted away from zero. For example, a Bayesian *p* value of .05 denotes that 95% of the resulting distribution is above (for positive relationships) or below (for negative relationships) zero. In acknowledgement of the large number of plausible values tests, we take a Bayesian *p* value <.01 as reliable. This method also allows the comparisons of two sets of plausible values relationships. For each relationship, we can take the differences between the averaged population correlation distributions. If the resulting difference distribution returns a Bayesian *p* value <.01, the difference in relationships is considered to be reliably greater than zero. This method was used to test whether relationships between SSRT_EXG3_ and the N1 are stronger than between SSRT_EXG3_ and P3.

### 2.5. Analysis Code and Output

The code used to analyse these data, as well as more detailed analysis output can be found in an Open Science Foundation repository at osf.io/rhktj. These materials include the code for EXG3 modelling of stop-signal task behaviour and the output of posterior predictive checks (Heathcote et al., 2018). The repository also includes the code analysing ERP waveforms, the output of ERP analysis at each electrode site of interest, the plausible values code and output for each site of interest.

## 3. Results

### 3.1. Stop-signal task behaviour

Of the 198 ± .85 stop trials (mean ± SEM), participants responded on average to 49.14% ± .45% of stop trials, indicating that the tracking algorithm was effective. Of the 479 ± 2.7 go trials, errors of omission and commission occurred on 2.99 ± .31 % and 7.41 ± 0.51% of trials, respectively. The mean stop-signal delay was 354.24 ± 9.63 ms (range = 89.42 – 705.29 ms). Mean RT on correct go trials was 585.56 ± 7.37 ms. Mean RT on stop-failure trials was faster than the average go RT (521.69 ± 6.09 ms). The slope of a regression of RT against trial number indicated that participants slowed their responses by an average of only 43.58 ± 7.63 ms between the start and end of the test session. Lastly, traditional estimation via the integration technique returned a mean SSRT_int_ of 196.91 ± 7.43 ms^5^.

### 3.2. Model selection and parameter estimation

Both models achieved convergence (i.e., 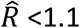). The model with trigger failure more closely matched the observed data than one without trigger failure and was selected by DIC by a wide margin (1685). Hence, we used the EXG3 model with trigger failure in all further analyses. Figure 1 shows a visualisation of the posterior distributions of the parameter estimates from this EXG3 model. The EXG3-based estimate of mean SSRT_EXG3_ had a mean of 132.35 ms with a 95% credible interval of [124.55, 140.17], which was around 65ms faster than SSRT_int_. Trigger failure rates were 22.27% [21.43, 24.82], indicating on average 22% of stop-failure trials were due to a failure to trigger an inhibitory response. Go failures occurred on 3.06% [2.83, 3.76] of go trials and the mean finishing time of the matching go runner was 596.51 ms [581.13, 611.84].

**Figure 1.**
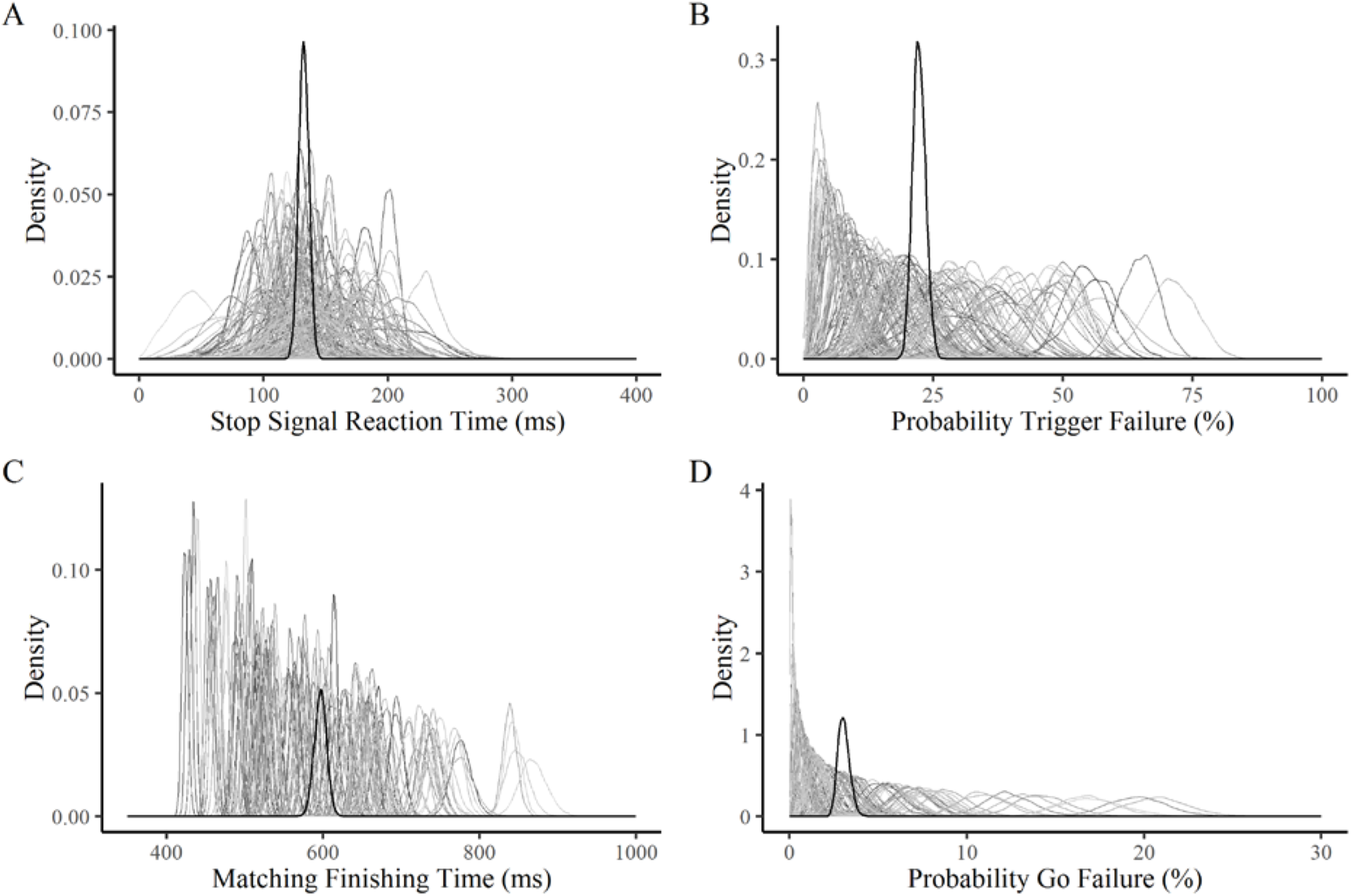
Posterior distributions of the individual (grey density lines) and group-level mean (black boldface density lines) parameters. Mean Stop-signal Reaction Time (ms; Panel A). Probability of Trigger Failure (Panel B). Mean finishing time for the matching go runner (ms; Panel C). Probability of Go Failure (Panel D). Each grey shaded distribution represents the posterior of a single participant. The black distribution represents the posterior distribution of the population-level mean parameter.

### 3.3. ERP summary

Table 1 shows summary statistics of ERP components. Figure 2 shows ERP waveforms and scalp topographies for each trial type and highlights that both N1 and P3 components were larger over the FCz electrode. The N1 peaked earlier and was larger for stop-success compared to stop-failure trials^6^. Similarly, the P3 emerged and peaked earlier, and was larger for stop-success than stop-failure trials.

**Figure 2.**
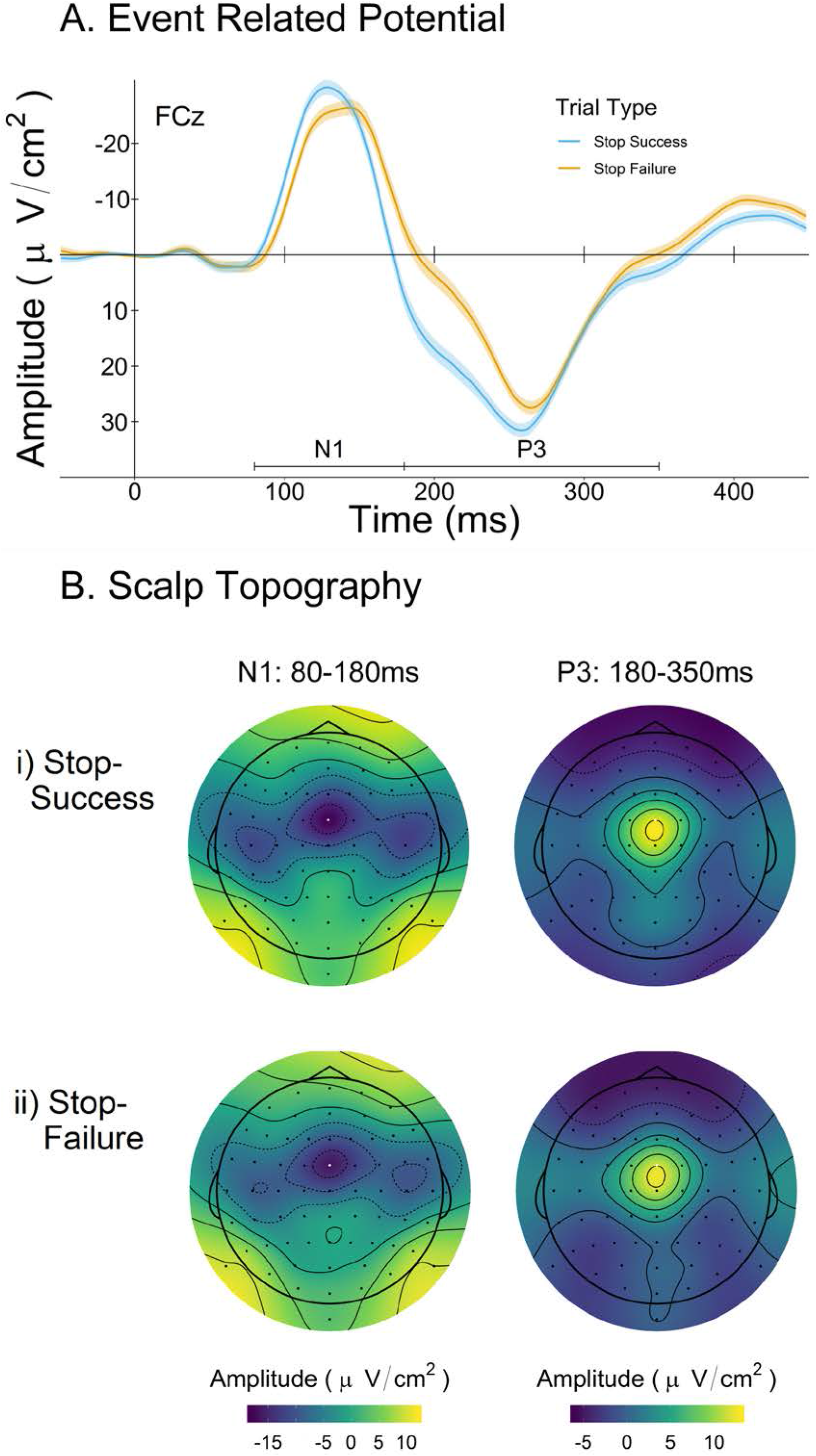
A. The stop-locked ERP waveform at FCz for stop-success and stop-failure trials, with lines indicating the measurement intervals for N1 and P3. B. Scalp topography of N1 and P3 on stop-success and stop-failure trials. Topographies display the average group amplitude over specified windows.

**Table 1.** Summary of the onset latency (ms), peak latency (ms), and amplitude (μv/cm^2^) of the N1 and P3 components measured at FCz and associated trial type differences.

| Measure | Stop Success<br>(SD) | Stop Failure<br>(SD) | Trial Difference<br>(BF<sub>10</sub>) | Effect<br>Size<br>($\delta$) |
| --- | --- | --- | --- | --- |
| **N1** |  |  |  |  |
| Onset Latency | 102.35 (13.87) | 105.34 (16.22) | 0.94 | -0.13 |
| Peak Latency | 132.56 (10.79) | 138.01 (12.64) | $6e^8$ | -0.02 |
| Peak<br>Amplitude | -33.61 (19.67) | -31.04 (21.06) | 8.67 | 0.17 |
| **P3** |  |  |  |  |
| Onset Latency | 220.35 (26.28) | 232.82 (24.96) | $2.13e^{10}$ | -0.52 |
| Peak Latency | 259.29 (22.27) | 267.81 (22.37) | $8.59e^5$ | -0.25 |
| Peak<br>Amplitude | 34.64 (22.91) | 30.21 (20.02) | $3.25e^3$ | 0.34 |

<sup>6</sup> Note that, in addition to the midline N1, topographical plots in Figure 2 show two weaker lateral foci were evident. As there was no discernible difference in the latencies of N1 between midline and lateral foci, we focussed on the midline N1 at FCz
*Note.* Effect sizes are for the success – failure trial type difference. Shading indicates at least positive evidence in favour of the trial type difference (i.e., $BF_{10} > 3$ ).
Note that larger N1 amplitude is represented by more negative values, whereas larger P3 amplitude is represented by more positive values. Likewise, for all ERP latency measures, a positive $r$ -value signifies a direct relationship (i.e., longer latency associated with longer RT and higher failure rate/slower SSRT). The same applies for $p$ -values derived from plausible value relationships.

Note that larger N1 amplitude is represented by more negative values, whereas larger P3 amplitude is represented by more positive values. Likewise, for all ERP latency measures, a positive *r*-value signifies a direct relationship (i.e., longer latency associated with longer RT and higher failure rate/slower SSRT). The same applies for ρ-values derived from plausible value relationships.

Table 2 provides summary statistics for the Pearson correlation analysis between SSRT_int_ and ERP component measures. Faster SSRT_int_ was associated with larger (i.e., more negative) N1 amplitude on both trial types. On stop-success trials, there was positive evidence in favour of a null relationship between N1 peak latency and SSRT_int_ (BF_10_ = .32) and equivocal evidence for a relationship between N1 onset latency and SSRT_int_ (BF_10_ = .35). In contrast, on stop-failure trials, N1 onset and peak latency were both associated with SSRT_int_. Paradoxically, delayed N1 onset and peak latency on stop-failure trials was associated with faster SSRT_int_.

**Table 2.**
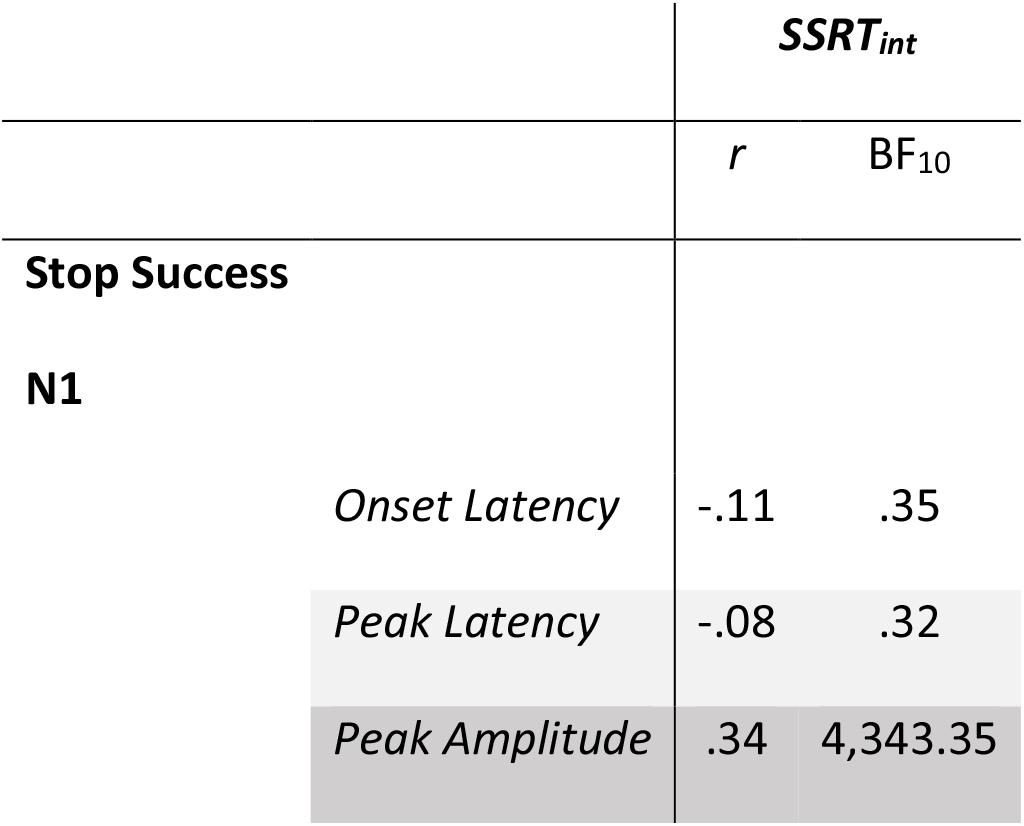
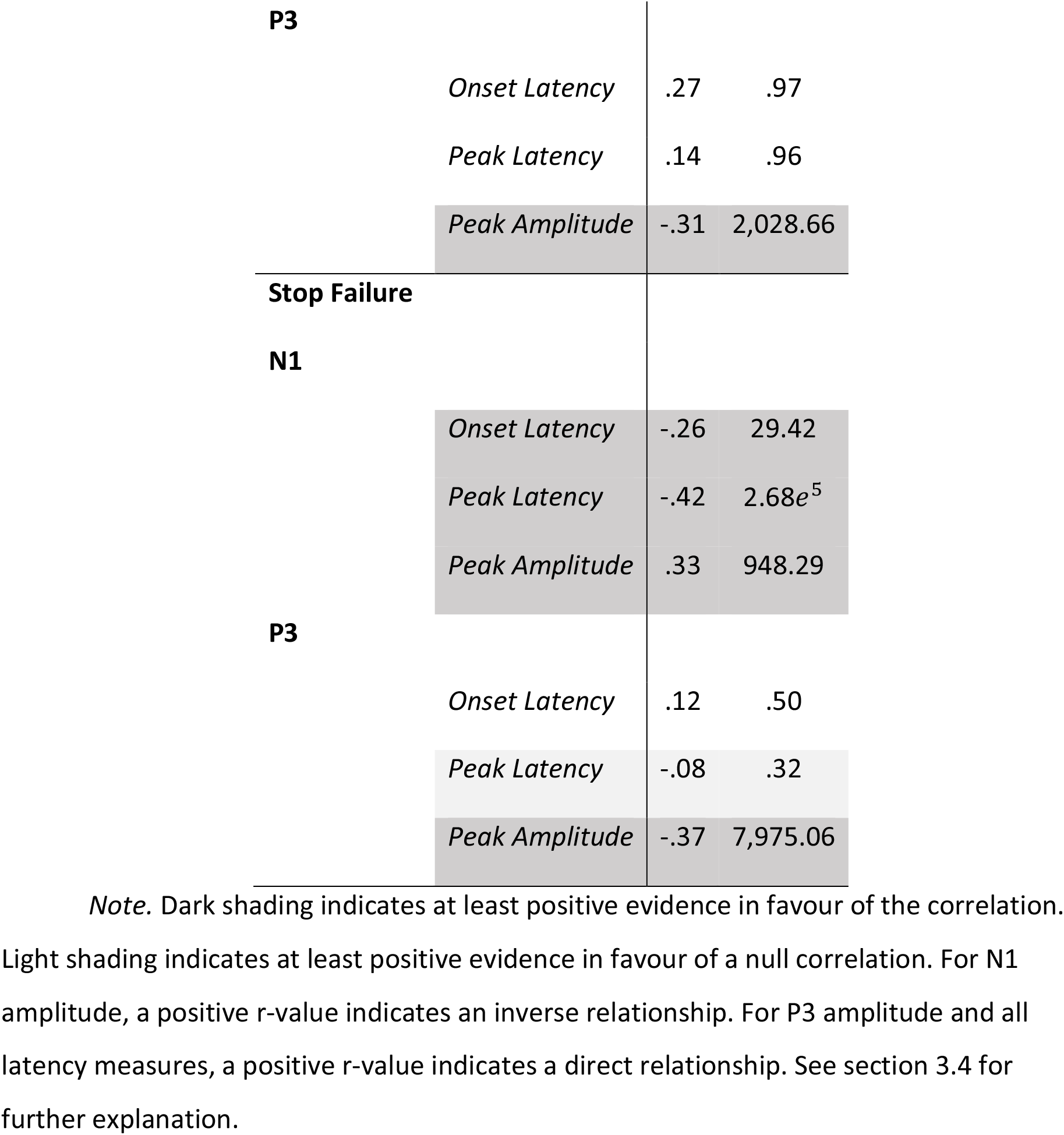
Pearson product moment correlations and associated Bayes factors for the relationships between SSRT_int_ and ERP component measures; onset latency (ms), peak latency (ms), and amplitude (μV/cm^2^).

|  |  | ***SSRT<sub>int</sub>*** |  |
| --- | --- | --- | --- |
| | | *r* | $BF_{10}$ |
| **Stop Success** |  |  |  |
| **N1** |  |  |  |
|  | *Onset Latency* | -.11 | .35 |
|  | *Peak Latency* | -.08 | .32 |
|  | *Peak Amplitude* | .34 | 4,343.35 |
| **P3** |  |  |  |
|  | *Onset Latency* | .27 | .97 |
|  | *Peak Latency* | .14 | .96 |
|  | *Peak Amplitude* | -.31 | 2,028.66 |
| **Stop Failure** |  |  |  |
| **N1** |  |  |  |
|  | *Onset Latency* | -.26 | 29.42 |
| | *Peak Latency* | -.42 | $2.68e^5$ |
|  | *Peak Amplitude* | .33 | 948.29 |
| **P3** |  |  |  |
|  | *Onset Latency* | .12 | .50 |
|  | *Peak Latency* | -.08 | .32 |
|  | *Peak Amplitude* | -.37 | 7,975.06 |

*Note.* Dark shading indicates at least positive evidence in favour of the correlation.
Light shading indicates at least positive evidence in favour of a null correlation. For N1 amplitude, a positive $r$ -value indicates an inverse relationship. For P3 amplitude and all latency measures, a positive $r$ -value indicates a direct relationship. See section 3.4 for further explanation.

### 3.5. Plausible values relationships with EXG3 model parameters

We first present the plausible values relationships between SSRT_int_ and EXG3 model parameters, referring to median ρ estimates, Bayesian *p* value, and 95% credible intervals. We then summarise the plausible values relationships between ERP component measures and EXG3 model parameters in Table 3. Small Bayesian *p*-values suggest that the distribution is shifted away from zero.

**Table 3.** Plausible value relationships. Median ρ, Bayesian p-values, and 95% credible intervals for the relationship between model estimated response inhibition parameters and ERP component measures; onset latency (ms), peak latency (ms), and amplitude (μv/cm^2^).

| Trial Type | Stop Signal Reaction Time |  |  | Probability of Trigger Failure |  |  | Matching Go Finishing Time |  |  |
| --- | --- | --- | --- | --- | --- | --- | --- | --- | --- |
| | $\rho$ | $p_{\text{Bayes}}$ | CI | $\rho$ | $p_{\text{Bayes}}$ | CI | $\rho$ | $p_{\text{Bayes}}$ | CI |
| **Stop Success** |  |  |  |  |  |  |  |  |  |
| **N1** |  |  |  |  |  |  |  |  |  |
| *Onset Latency* | .063 | .242 | [-.11, .24] | .141 | .047 | [-.02, .3] | .011 | .445 | [-.15, .17] |
| *Peak Latency* | -.004 | .502 | [-.18, .17] | -.055 | .276 | [-.22, .11] | -.002 | .508 | [-.16, .16] |
| *Peak Amplitude* | .295 | <.001 | [.12, .45] | .253 | <.001 | [.09, .4] | -.103 | .111 | [-.26, .05] |
| **P3** |  |  |  |  |  |  |  |  |  |
| *Onset Latency* | .191 | .026 | [0, .37] | .11 | .116 | [-.07, .28] | -.198 | .014 | [-.36, -.03] |
| *Peak Latency* | .174 | .026 | [0, .34] | .099 | .119 | [-.07, .26] | -.141 | .045 | [-.29, .02] |
| *Peak Amplitude* | -.348 | <.001 | [-.49, -.18] | -.219 | .005 | [-.37, -.06] | .303 | <.001 | [.15, .44] |
| **Stop Failure** |  |  |  |  |  |  |  |  |  |
| **N1** |  |  |  |  |  |  |  |  |  |
| *Onset Latency* | -.162 | .04 | [-.33, .01] | -.206 | .008 | [-.36, -.04] | .145 | .036 | [-.01, .3] |
| *Peak Latency* | -.226 | .007 | [-.39, -.05] | -.374 | <.001 | [-.51, -.22] | .184 | .011 | [.03, .33] |
| *Peak Amplitude* | .268 | <.001 | [.1, .42] | .208 | .006 | [.05, .36] | -.129 | .061 | [-.28, .03] |
| **P3** |  |  |  |  |  |  |  |  |  |
| *Onset Latency* | .173 | .035 | [-.01, .35] | .031 | .363 | [-.14, .2] | -.148 | .045 | [-.31, .02] |
| *Peak Latency* | .05 | .289 | [-.13, .22] | -.119 | .087 | [-.28, .05] | -.039 | .334 | [-.19, .12] |
| *Peak Amplitude* | -.37 | <.001 | [-.51, -.2] | -.243 | .002 | [-.39, -.08] | .338 | <.001 | [.19, .47] |

*Note.* $\rho$ = Median of the posterior distribution of the population correlation. $p_{\text{Bayes}}$ = Bayesian p value. CI = Credible Interval [2.5%, 97.5%]. Shading indicates reliable relationships (i.e., $p_{\text{Bayes}} < .01$ ). For N1 amplitude, a positive p-value indicates an inverse relationship. For P3 amplitude and all latency measures, a positive p-value indicates a direct relationship. See section 3.4 for further explanation.

#### 3.5.1. Relationships between traditional SSRT estimates and EXG3 model parameters

SSRT_EXG3_ was faster than SSRT_int_ (132.35 vs 196.91 ms), and the two values were moderately correlated, ρ = .537, *p*_Bayes_ < .001 [95% CI: .373, .670). SSRT_int_ was strongly positively correlated with trigger failure rate, ρ = .846, *p*_Bayes_ < .001 [.787, .89], but only weakly with go failure rate, ρ = .246, *p*_Bayes_ = .001 [.088, .391]. The difference of the plausible values distributions of trigger failure with SSRT_int_ and SSRT_EXG3_ had a Bayesian *p*-value = .002, showing that only ≈.2% of the difference distribution overlaps zero. This indicates that trigger failure was a better predictor of traditional SSRT than the EXG3 estimation of SSRT.

#### 3.5.2. Relationships between ERP measures and SSRT_EXG3_

As found with SSRT_int_, faster SSRT_EXG3_ was associated with a delayed N1 peak latency on stop-failure trials. However, the relationship with N1 onset latency was equivocal (Table 3). For both trial types, faster SSRT_EXG3_ was associated with larger N1 and P3 peak amplitude (Table 3), and there was no reliable difference between the plausible values distributions of N1 and P3 amplitude for either trial types (*p*_Bayes_ >.01). This does not support our prediction that relationship with SSRT_EXG3_ would be stronger with N1 than with P3. Overall, there was no reliable relationship between the variability of SSRT_EXG3_ and any ERP component measures (see osf.io/rhktj).

#### 3.5.3. Relationships between ERP measures and Probability of Trigger Failure

Trigger failure showed an identical pattern of correlations with ERP measures as SSRT_int_. Higher rate of trigger failure was associated with earlier N1 onset and peak latency measures for stop-failure trials only (Table 3). In addition, for both trial types, higher rate of trigger failure was associated with smaller N1 amplitude, suggesting that lower attention to the stop-signal increases the rate of trigger failure. The rate of trigger failure was not associated with P3 onset/peak latency for either trial type. However, for both trial types, higher trigger failure rate was associated with smaller P3 amplitude (average ρ ≈ −.23; Table 3).

#### 3.5.4. Relationships between ERP measures and Finishing Time of the Matching Go Runner

On both trial types, slower finishing time of the matching go runner was associated with larger P3 amplitude (average ρ ≈ −.32; Table 3).

#### 3.5.5. Relationships between ERP measures and Probability of Go Failure

On stop-failure trials, higher go failure rate was associated with earlier N1 latency, ρ = −.221, *p*_Bayes_ = .004 [.37, −.06]. No other relationships were reliable.

### 3.6. N1 peak latency and paradoxical relationships with TF and SSRT

We ran exploratory analyses to understand the paradoxical finding that faster N1 latency on stop-failure trials is associated with both higher rates of trigger failure and slower SSRT. We examined whether there are differences in the pattern of ERP effects as a function of performance by extracting ERP waveforms for three groups of participants based on a tertile split of the median of the participant-level posteriors.

For trigger failure, Figure 1.B shows that the distribution of trigger failure had a strong right skew that spread out past ≈20%. The tertile split resulted in three groups with Low (<14%), Mid (>= 14% and =<25.8%) and High (>25.8%) trigger failure (n=52 per group). The groups had similar age (Low = 20.4; Mid = 21.5; High = 20.65) and gender (Female: Low = 48.1%; Mid = 53.8%; High = 57.7%) distributions.

Likewise, the tertile split of the participant-level posterior SSRT_EXG3_ values resulted in three groups with Fast (<124ms), Mid (>= 124ms and =<150ms) and Slow (>150ms) SSRT_EXG3_ (n=52 per group) with similar age (Fast = 20.3, Mid = 20.9, Slow = 21.4 years) and gender (Female: Fast = 48.1%; Mid = 57.7%; Slow = 53.8%) distributions. Twenty-seven (51.9%) participants in the Low Trigger Failure group were also in the Fast SSRT group, and 28 (53.8%) participants in the High Trigger Failure group were in the Slow SSRT group. However, the overlap for the two Mid groups was smaller with only 19 participants (36.5%).

ERPs for each level of the Trigger Failure and SSRT groups are shown in Figure 3 and summary statistics for N1 peak latency are shown in Table 4. N1 peak latency was analysed using a Bayesian mixed ANOVA with default priors (JASP Team, 2018) and two factors: group (3 performance group levels) and trial type (stop-failure, stop-success). For trigger failure, the performance group levels were High, Mid, Low, and for SSRT they were Slow, Mid, and Fast.

**Figure 3.**
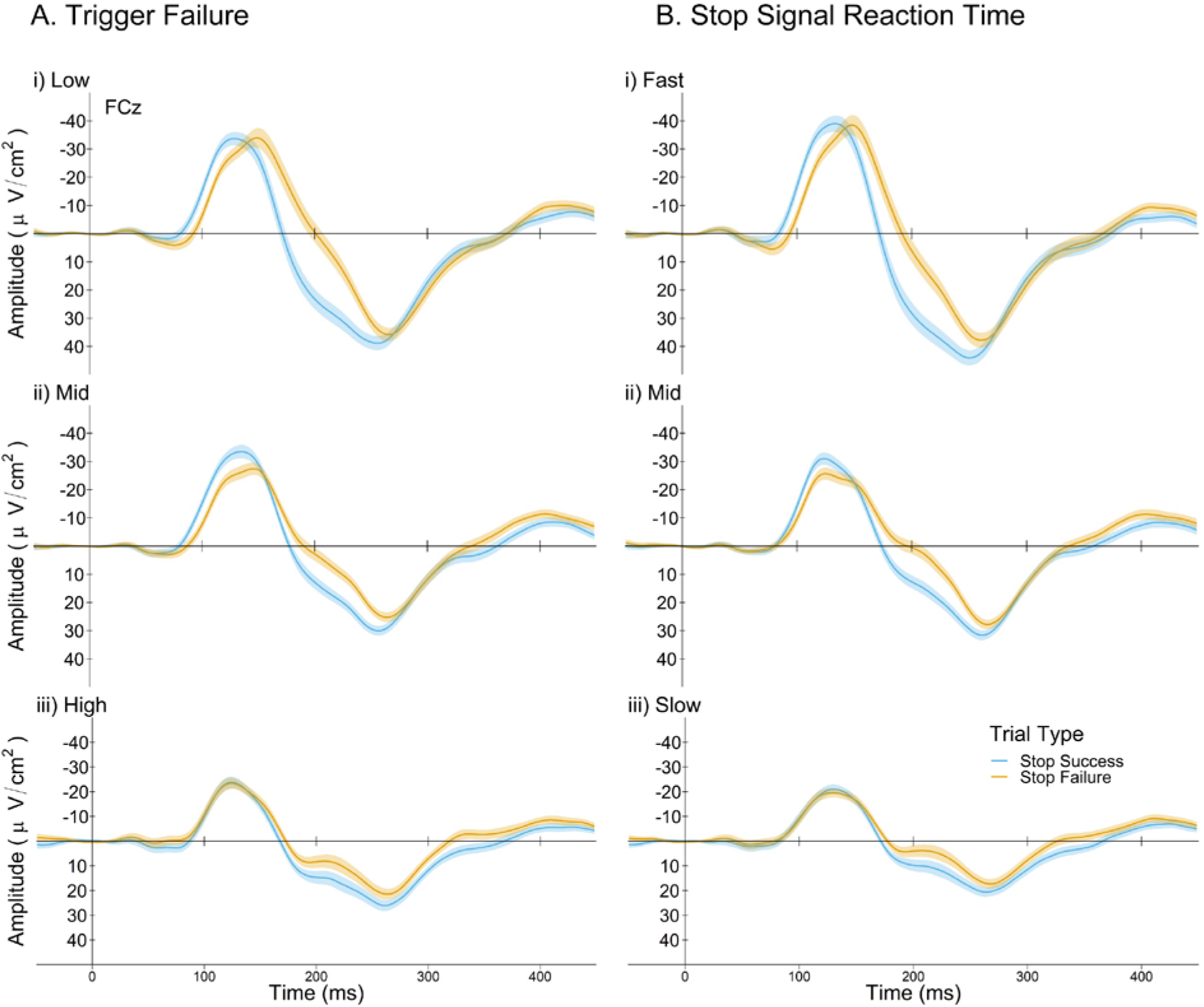
Stop-locked ERPs at FCz for stop-success (blue) and stop-failure (orange) trials for Trigger Failure (A) and SSRT (B) groups. Shaded areas represent within subject confidence intervals.

**Table 4.**
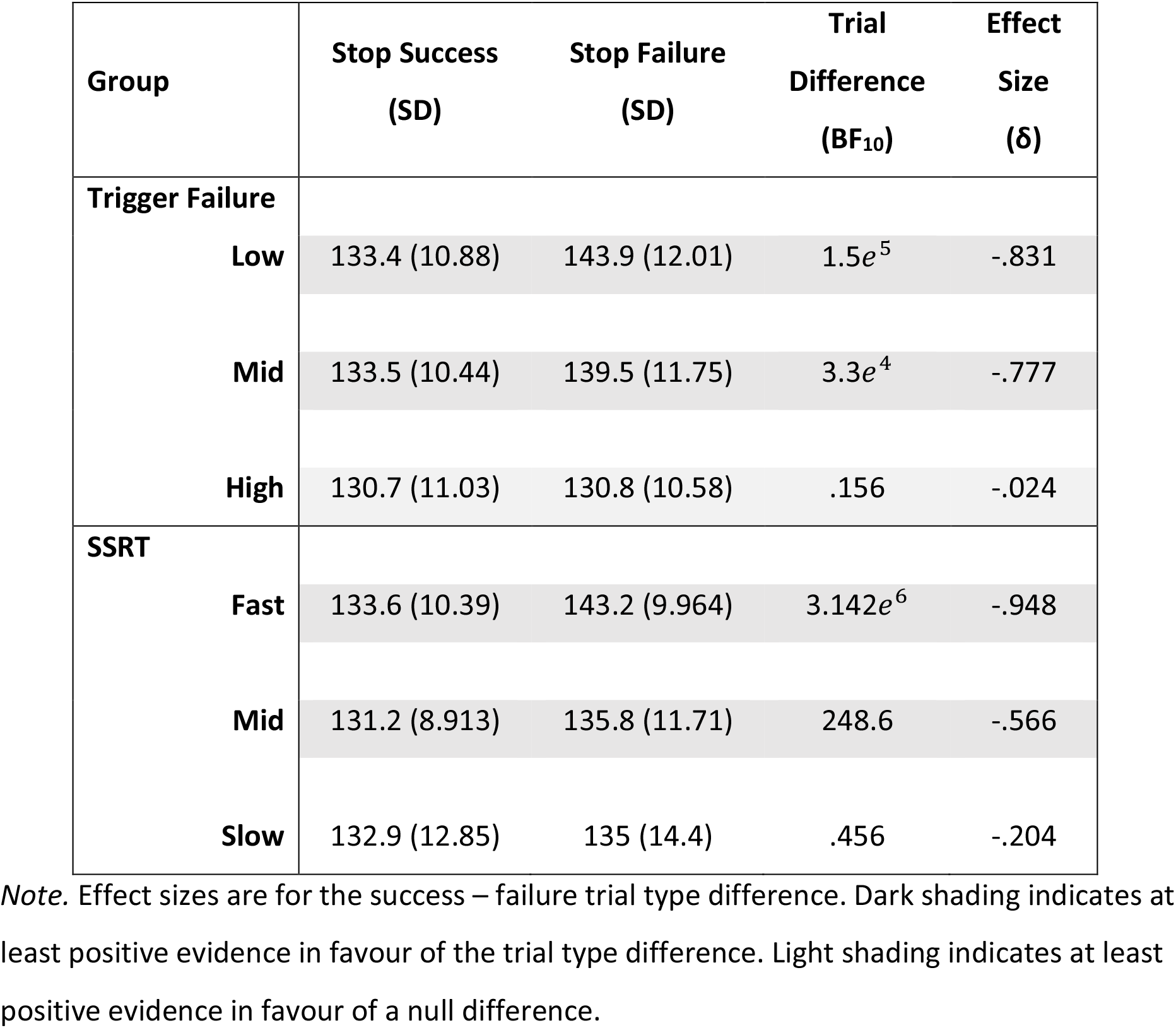
N1 peak latency trial type differences between tertile splits of Trigger Failure and SSRT.

Analysis of trigger failure groups resulted in very strong evidence for the model containing the two main effects (BF_10_ = 3.62*e*^9^). The interaction model was also supported when controlling for the main effects of group and trial type (BF Inclusion = 3.1*e*^4^). For Low and Mid (but not High) trigger failure groups, the N1 peaked earlier for stop-success than stop-failure trials (Table 4). Analysis of the SSRT groups also showed very strong evidence for a model containing the two main effects (BF_10_ =7.9*e*^7^) and strong evidence for the interaction model, after controlling for both main effects (BF Inclusion = 36.833). For both the Fast and Mid SSRT groups, N1 peaked earlier for stop-success than stop-failure trials, but this evidence for this effect was equivocal in the Slow group (Table 4).

To further clarify these interactions, we examined differences in N1 peak latency across groups for each trial type. On stop-success trials, there were no group differences. On stop-failure trials, N1 peak latency was earlier for the High trigger failure than either the Mid or the Low trigger failure groups (BF_10_ = 159.06, δ = .726; BF_10_ = 2.4*e*^5^, δ = 1.099), but there was only equivocal evidence for a difference between Low and Mid trigger failure groups (BF_10_ = 1.001, δ = .332). Similarly, N1 peaked earlier for the Fast SSRT than either the Mid or the Slow groups (BF_10_ = 33.317, δ = .621; BF_10_ = 27.514, δ = .604), but the evidence favoured a null difference between Mid and Slow SSRT groups (BF_10_ = .218, δ = .05).

In sum, while N1 peaked earlier for stop-success than stop-failure trials in mid- and high-performing participants, poorer performing participants (i.e., High trigger failure; Slow SSRT) showed no differentiation in N1 peak latency between the two trial types. In fact, for stop-failure trials, the N1 peaked earlier in poor relative to good performers. Thus, the earlier N1 peak for stop-failure trials in participants with high trigger failure and slow SSRT appears to drive the relationship between N1 peak latency and both trigger failure rate and SSRT in the whole group analyses.

Finally, we examined whether SSRT and trigger failure groupings differed in N1 amplitude and found no support for an interaction between trial type and either the trigger failure (BF_Inclusion_ = 1.325) or the SSRT (BF_Inclusion_ = .073) grouping^7^. In sum, strong differences in N1 latency but not amplitude were shown between performance groupings based on stop-signal task performance.

## 4. Discussion

This is the first study to comprehensively examine the relationship between stop signal ERPs and behavioural parameters derived from a parametric model of response inhibition that accounts for failures of both stop and go processes. Trigger failure was non-negligible and substantially influenced the interpretation of ERP components associated with response inhibition during the stop-signal task. Individual differences in both SSRT and trigger failure rate yielded novel insights into the cognitive and neural processes that underlie response inhibition^8^. Below, we first compare the model-based estimates of SSRT with traditional non-parametric estimates, then discuss the interpretations for each ERP component, and the implications for response inhibition models.

### 4.1. Non-parametric vs. parametric estimates of SSRT

Consistent with recent studies, this parametric estimation technique resulted in an attenuation of SSRT compared to the traditional non-parametric integration method (Band et al., 2003; Jana, Hannah, Muralidharan, & Aron, 2020; Matzke, Love, et al., 2017; Skippen et al., 2019; Weigard, Heathcote, Matzke, & Huang-Pollock, 2019). Importantly, SSRT_int_ was only moderately correlated with SSRT_EXG3_, but was strongly associated with trigger failure. These findings support the argument that traditional SSRT estimates are biased by the assumption that trigger failure rates are negligible. As a result, traditional SSRT estimates confound the speed of the inhibition process (i.e., SSRT) with the reliability of the inhibition process (i.e., trigger failure).

### 4.2. The stop-N1 – an index of automatic inhibition processes?

Consistent with previous studies using an auditory stop-signal task, N1 amplitude was larger for stop-success than stop-failure trials (Bekker, Kenemans, et al., 2005; Hughes et al., 2012; Kenemans, 2015; but see Ramautar, Kok, & Ridderinkhof, 2006). This ‘stop-N1’ effect suggests greater attentional allocation to the stop signal on trials that eventually lead to successful inhibition. Moreover, faster SSRT (both SSRT_int_ and SSRT_EXG3_) was moderately correlated with increased N1 amplitude for both stop-success and stop-failure trials^9^. These findings are consistent with a tonically active, top-down potentiation of the impact of the stop signal (Kenemans, 2015). Specifically, Kenemans has proposed that the right inferior frontal gyrus (rIFG) potentiates the link between the detection of the stop signal and an automatic, speeded inhibitory response. Increased attentional allocation, reflected by the ‘stop-N1’ effect, appears to modulate the probability of successful inhibition and result in a faster, automatically driven, stop process.

The notion of an automatic response inhibition process is supported by both the fast estimates of SSRT (≈130 ms; see also Skippen et al., 2019; Matzke, Hughes, et al., 2017) and by motor evoked potential studies that suggest the inhibition processes occurs around 140ms (Coxon, Stinear, & Byblow, 2006; Jana, Hannah, Muralidharan, & Aron, 2020; Raud & Huster, 2017; Waldvogel et al., 2000; Wilcoxon, Nadolski, Samarut, Chassande, & Redei, 2007). In fact, a recent multi-modal study by Jana et al. (2020) showed that the cascade of processing that occurs during the stop-signal task suppresses muscle activity within 160 ms after a visual stop-signal onset. As the registration of auditory signals occurs around 30 ms earlier than visual signals (Ramautar, Kok, & Ridderinkhof, 2006; Jain, Bansal, Kumar, & Singh, 2015), the 130 ms estimate of SSRT found here is consistent with the fast inhibition process described by Jana et al. (2020).

Importantly, we present the first evidence that a ‘stop-N1’ effect is present, not only for its amplitude, but also for its latency. In better performing participants, N1 emerged and peaked earlier on stop-success than stop-failure trials. This ‘stop-N1 latency’ effect is consistent with the horse-race model: earlier processing of the stop-signal allows relatively more time for the stop processes to win the race. Just as shorter stop-signal delays are easier to inhibit, faster processing of the stop-signal (i.e., earlier N1 latency) leads to easier inhibition. Tentatively, we suggest that both increased (i.e., stop-N1 amplitude) and earlier (i.e., stop-N1 latency) attentional allocation modulates the probability of successful inhibition. The finding that N1 peaks earlier and is larger on stop-success than stop-failure trials suggests that it indexes a cognitive process that moderates the inhibition process.

Consistent with this conclusion, participants with ADHD do not show a ‘stop-N1’ amplitude effect (Bekker, Overtoom, et al., 2005; Overtoom et al., 2009) and have both slower SSRT and higher trigger failure rates compared to controls (e.g., Weigard, et al., 2019). Weigard et al. (2019) recently showed that trigger failure is more sensitive than SSRT to differences between ADHD and matched controls. While the relationship between trigger failure and N1 effects has not been investigated in ADHD, Matzke, Hughes, et al. (2017) showed that people with schizophrenia had higher trigger failure rate and slower SSRT than controls. In the schizophrenia group, increased rate of trigger failure was correlated with earlier N1 peak latency.

We replicated the negative correlation between N1 peak latency and trigger failure in a large sample of healthy participants who produce a wide range of trigger failure rates and showed that this correlation is only present on stop-failure trials. Matzke, Hughes et al. (2017) suggested that the somewhat paradoxical relationship between N1 peak latency and trigger failure may result from reduced amplitude of a late sub-component of the N1. However, as N1 onset latency was also negatively correlated with trigger failure, it is unlikely that the late N1 sub-component can completely explain the negative correlation between N1 peak latency and trigger failure.

Alternatively, we suggest that poor performance in the stop-signal task is mediated, at least partly, by a lack of connection between activation of the stop process and the outcome of the trial (e.g., Kenemans, 2015). In previous literature, failure to utilise the automatic inhibitory pathway is thought to be indicated by the absence of a ‘stop-N1’ amplitude effect (see Bekker, Overtoom, et al., 2005; Overtoom et al., 2009). Our findings show that poor performance is more strongly reflected by the absence of a ‘stop-N1’ latency effect. We speculate that modulation of the N1 more generally may reflect the utilisation of the automatic inhibitory pathway. Further investigation is necessary to understand whether N1 amplitude and latency differentially affect stop-signal task performance.

Interestingly, the N1 component showed similar timing as an EEG beta frequency response that occurs ≈150ms after the stop signal (i.e., well before traditional estimates of SSRT) and has also been associated with successful stopping (Huster, Schneider, Lavallee, Enriquez-Geppert, & Herrmann, 2017; Swann et al., 2009; Swann et al., 2012; Wagner, Wessel, Ghahremani, & Aron, 2018; Wessel, Conner, Aron, & Tandon, 2013). This beta effect shows similar trial type differences as both the N1 and the P3 but is temporally aligned with the N1 and has a rIFG source (Wagner, Wessel, Gharahmeni, & Aron, 2018). More work is needed to examine whether the N1 and beta effects represent the same automatic inhibition mechanism associated with tonic rIFG activation (Kenemans, 2015).

### 4.4. Manifestation of Response Inhibition and the P3

In contrast to the hypothesis that P3 represents the electrophysiological manifestation of the stop process, we found no evidence for a relationship between P3 peak or onset latency and SSRT. In a recent meta-analysis of the association between SSRT and the P3, Huster, Messel, Thunberg, and Raud (2019) found a correlation coefficient of *r* = 0.41 (95% confidence interval = 0.02 to 0.69)^10^ between P3 onset latency and SSRT, despite the fact that P3 onset latency was often as much as 100 ms later than the SSRT estimate. Therefore, the similar latency difference between P3 and SSRT_EXG3_ in our data is unlikely to explain the absence of a P3 and SSRT relationship.

In an empirical sample, Huster et al. (2019) showed that SSRT correlated with both N2 and P3 peak latency from a visual stop-signal paradigm. They also showed that P3 onset latency correlated with both go RT and the probability of stopping. They suggested that the P3 latency generally reflects a type of speed-accuracy trade-off process that balances the different demands of go and stop stimuli in the stop-signal task. While we cannot speak to the N2 effects using an auditory paradigm, we report no relationship between P3 onset latency and go RT to support this interpretation. Instead, we suggest the P3 represents an evaluation process, which follows the success or failure of the inhibitory process. This explanation allows for the large temporal lag between SSRT and P3 onset latency and is still in line with studies that find significant correlations between SSRT and both the peak and onset latency of the P3. For example, the earlier P3 onset latency for stop-success trials suggests that the evaluation process starts earlier when there is not a concurrent error detection process like that found in stop-failure trials.

Alternatively, Wessel and colleagues have recently shown evidence that the stop P3 shares common neural generators with the P3 elicited in tasks involving infrequent stimulus detection (Waller, Hazeltine, & Wessel, 2019; Wessel & Huber, 2019). If the P3 indexes a stimulus detection process (or the evaluation of stimulus detection), the larger P3 amplitude on stop-success compared to stop-failure trials might index the successful detection of the stop-signal on the former and/or poor detection on the latter trial type. This interpretation is consistent with the horse-race model, as well as with our interpretation of the N1 effect as an early index of inhibitory success. Speculatively, we suggest that overlapping attentional processes are required to both detect the novelty of the stop-signal and trigger an inhibitory response and may explain the otherwise unexpected relationship between trigger failure and P3 amplitude across trial types.

There is still much debate about the role of the fronto-central P3 in response inhibition. While our findings do not rule out the P3’s role in response inhibition, we show that early, attentional processes reflected in N1 peak latency are strongly related to the success or failure of response inhibition, and that these processes are more strongly related to individual variability in stop-failure than the P3. Future studies need to investigate the relative contribution of these two components to inhibitory control ability.

### 4.5. Caveats and future work

Consistent with many earlier stop-signal studies, we used an analysis approach that maximised automated pre-processing of EEG data, while adhering to conventional processing guidelines (e.g., Lansbergen et al., 2007; González-Villar, Bonilla & Carrillo-de-la-Peña, 2016; Senderecka, Grabowska, Szewczyk, Gerc & Chmylak, 2012; Dimoska & Johnstone, 2008), rather than more recently used independent component analyses (ICA) methods (e.g., Wessel & Aron, 2015). Huster et al. (2019) showed no interpretable differences in the size of the correlations between SSRT and ERP components derived from ICA-based vs. conventional artefact removal pipelines. Huster et al. (2019) also showed that the relationship between SSRT and P3 onset latency was similar whether using a 50% peak amplitude estimation (used here) or the approach used by Wessel and Aron (2015). We also re-estimated P3 onset latency using a 10% fractional area latency (e.g., Raud & Huster, 2017) and again found no evidence for a reliable relationship between stop-success P3 onset latency and SSRT_int_ or SSRT_EXG3_. Therefore, we do not believe that either pre-processing or component measurement differences between studies had considerable influence on our findings or interpretations.

As discussed in Skippen et al. (2019), the current paradigm utilised a more demanding go task (a number parity task) than typical stop-signal paradigms (e.g., X vs. O discrimination). While this may have led to relatively high trigger failure rates in some participants, it is this variability in trigger failure that permitted analysis of individual differences in response inhibition parameters and their relationship with stop-trial ERPs. Previous studies examining trigger failures in healthy participants have low trigger failure rates (≈ 9%), but also low inter-individual variability and substantial ceiling effects (e.g., Matzke, Hughes, et al., 2017). While trigger failure and go failure parameters are still relatively novel in the stop-signal task literature, the EXG3 model can be readily applied to existing data to test the strength of our interpretation of ERP components of response inhibition. Differences between paradigms provide an opportunity to better understand trigger failure, SSRT, and response inhibition.

## 5. Conclusion

Contemporary models of response inhibition are challenging the notion that mean SSRT can fully capture individual differences in inhibition ability. Novel parameters, like trigger failure and go failure, together with SSRT estimates that consider trial-by-trial variability, allow richer characterisation of inter- and intra-individual variability in stop-signal task behaviour. We found this more detailed approach to characterising the processes involved in response inhibition provided evidence that the early, attention-related N1 is sensitive to individual differences in both the speed and the reliability of the inhibition process. Furthermore, modulations of N1 latency in participants with slow SSRT and/or high trigger failure rates indicated that, poor inhibitors exhibit a missing link between attending to the stop-signal and inhibition success. Further work is needed to extend our understanding of these N1 modulations to clinical groups with poor inhibitory control (i.e., ADHD, schizophrenia) to gain insight into the underlying causes of inhibitory deficits. Moreover, further investigation of the connection between these novel response inhibition parameters and rIFG activation (which is affected in ADHD and schizophrenia patients) could provide valuable insights into current models of inhibitory control. In conclusion, attentional mechanisms involved in eliciting the inhibition process appear to be just as, or possibly even more, important as the speed of the inhibition process in explaining differences in response inhibition.

## Acknowledgements

We thank Gavin Cooper for paradigm programming and members of the Age-ility Project for assistance with data collection/entry. Special thanks to Montana McKewen and Patrick Cooper for their contributions to project management. We would also like to thank participants for their time.

## Footnotes

1 Non-parametric estimation techniques make no assumptions about the form of the distributions of the go or stop runners (see Logan & Cowan, 1984; Matzke, Verbruggen, & Logan, 2018).

2 This study used the BEESTS procedure with only trigger failure and did not estimate go failure.

3 This was assessed by regressing RT on trial number and then using the slope of the fit to estimate slowing over the course of the task.

4 We also attempted to implement ADJAR level-2 (see Bekker, Kenemans, et al., 2005) but data did not converge for a number of participants.

5 It should be noted that these results are based upon the final sample of participants, who were screened on several behavioural criteria, including the response rate to stop trials, omission and commission errors, and slowing of go RT (see *2.3.1*).

6 Note that, in addition to the midline N1, topographical plots in Figure 2 show two weaker lateral foci were evident. As there was no discernible difference in the latencies of N1 between midline and lateral foci, we focussed on the midline N1 at FCz

7 Following a reviewer’s comment that the peak amplitude measure is taken at different latencies for each trial type, we repeated the N1 peak amplitude analyses using an amplitude measure which is consistent across both trial types. Firstly, we took the ERP to stop trials, averaged across trial types for each participant, and estimated the peak latency of the N1. Peak amplitude for each trial type was then calculated as described in *2.3.3. Electrophysiological Data*. There was no evidence to support an interaction between SSRT performance grouping and trial type on N1 amplitude (BF_Inclusion_ = .157). However, the interaction model was supported between trigger failure group and stop trial type (BF_Inclusion_ =3.53). Further testing revealed that ‘stop-N1’ effects were present for the mid trigger failure group (BF_10_ = 1322, δ = −0.635), but not the low (BF_10_ = .33, δ = - 0.167) or high (BF_10_ = .152, δ = −.016).

8 A reviewer cited an unpublished study (osf.io/kpa65) reporting graphical evidence for substantial violations of the context independence assumption made by all parametric and non-parametric SSRT estimates in a large number of studies, although the reliability of these violations were not checked statistically. The paper argues that such violations may be misidentified as trigger failures. We applied the same graphical check to our data and found no evidence of such violations (see osf.io/rhktj).

9 This finding has been reported before in a visual stop-signal paradigm with a collapsed sample of schizophrenia patients and matched controls based on non-parametric SSRT estimates (Hoptman et al., 2018).

10 Note that the authors of this meta-analysis show that the average power of the published studies to detect an effect was below 50% for one-sided tests.

